# Profile of the B cell receptor repertoire and antibody responses upon 17DD-YF vaccine boosting

**DOI:** 10.1101/2025.06.24.660700

**Authors:** Christina A. Martins, Milene Barbosa Carvalho, João Diniz Gervásio, Carlena Navas, Luciana Zuccherato, Marcele Rocha, Manuela Emiliano Ferreira, Camila Souza, Adriana de Souza Azevedo Soares, Brenda de Moura Dias, Nathalia dos Santos Alves, Sheila Maria Barbosa de Lima, Waleska Dias Schwarcz, Andréa Marques Vieira da Silva, Ana Paula Dinis Ano Bom, Camilla Bayma Fernandes, Renata Carvalho Pereira, Mauro M. Teixeira, Jason Lavinder, Gregory Ippolito, Liza Figueiredo Felicori

**Affiliations:** Departamento de Bioquímica e Imunologia, Instituto Ciências Biológicas, Universidade Federal de Minas Gerais, Belo Horizonte, Minas Gerais, Brazil; Departamento de Ciência da Computação, Universidade Federal de São João del Rei, São João del Rei, Minas Gerais, Brazil; Departamento Experimental e Pré-Clínico, Instituto de Tecnologia em Imunobiológicos Bio-Manguinhos, Fiocruz, Rio de Janeiro, Brazil; Department of Molecular Biosciences, The University of Texas at Austin, Austin, Texas, United States

**Keywords:** *Flavivirus* Immunity, Immunoglobulin Repertoire, Antibody Response, B Cell Receptor

## Abstract

Although the 17DD-YF vaccine is highly effective in conferring long-term immunity, recent studies suggest that booster doses may be necessary in endemic areas. In this study, we characterized the dynamics of the B cell repertoire following 17DD-YF booster vaccination and examined its impact on antibody kinetics. Using a combined approach of serological analysis and B cell receptor (BCR) sequencing, we investigated molecular features of the antibody response in four vaccinated individuals from a YFV-endemic region.

Our results indicate that the booster induces a transient antibody response, with limited effect on the neutralization capacity against DENV1-4 and ZIKV. Notably, we observed a rapid clonal expansion of B cells within 14 days post-vaccination, predominantly involving IgA and IgG isotypes. However, this expansion was short-lived, with reduced magnitude and persistence of these lineages by day 28. Interestingly, some of the expanded lineages shared sequence similarity with previously characterized anti-DENV and anti-ZIKV antibodies, although further validation is required to assess their functional relevance and presence in circulating serum.

Together, these findings highlight the transient nature and isotype composition of the antibody response following YFV revaccination and provide insights into the complex interplay between pre-existing immunity and recall responses in individuals living in endemic regions.

## 1. Introduction

*Yellow fever virus* (YFV) is an important human pathogen that is transmitted by infected mosquitoes of the genera *Aedes* and *Haemagogus*. It is endemic in tropical regions of Sub-Saharan Africa and South America and belongs to the genus *Flavivirus* that comprise more than thirty antigenically related viruses, including Dengue (DENV) and Zika (ZIKV) [1–2]. Infections caused by YFV may vary from asymptomatic to severe illness, which represents a high risk of morbidity and mortality due to the occurrence of haemorrhagic manifestations and hepatorenal syndromes [3–4]. To date, there is no specific treatment for YFV disease and vector control is challenging. The recent YFV outbreaks that happened in Brazil (2016-2019) have resulted in mortality 1.5 times higher compared to past epidemics that occurred in the country, revealing that YFV remains a persistent threat despite the availability of an effective vaccine and good vaccination coverage [5–6].

The YFV vaccine (17DD, 17D, 17D-213) is live-attenuated and is recognized as one of the best vaccines ever developed. A single dose of the YFV vaccine confers prolonged immunity lasting multiple decades and the vaccine has an excellent safety profile [7–8]. However, studies about the duration of the YFV vaccine immunity have shown a time-dependent loss of the YFV-neutralizing antibodies (nAb) titers 10 years after the vaccination, and the restauration of this humoral immunity with revaccination [9–11]. Therefore, booster regimens may be needed to maintain effective protection and could benefit specific groups of populations, especially those living in endemic areas with high risk for virus transmission [11].

It is well established that after vaccination, YFV-specific memory B cells are observed as early as 14 days and peak between three and six months later [6, 12]. nAbs are detected in the serum within a few days following vaccination [6, 13–15] and that serum nAbs can persist for up to 60 years [8, 16]. Most of these nAbs target the envelope protein (E) [17–18], which is the main structural component of the flaviviruses. The DII and DIII domains of the E protein, including a highly conserved epitope located at the tip of the DII domain that includes the fusion loop (FL), are some of the antigenic sites recognized by the YFV-nAbs [12, 19]. Given the fact that flaviviruses share a high degree of structural identity, most of these epitopes are conserved, and some of the nAbs may cross-neutralize even distantly related viruses [20–24].

The current knowledge of the human antibody repertoire elicited after the YFV vaccination remains limited to two studies, the first, compared the antibody response between young and middle-age individuals after vaccination [25], while the second characterized hundreds of YFV E-specific mAbs from two flavivirus-naïve donors who were vaccinated for the first time with the 17D-YF vaccine [12]. Although these studies provided important insights into the B cell dynamics and antibody cross-reactivity related to the 17D-YF vaccine response in *Flavivirus* unexperienced individuals [12], it still lacks a deep characterization of the features of the antibody repertoire induced by 17DD-YF vaccine in the context of revaccination of a *Flavivirus* endemic cohort.

To address this, we longitudinally characterized the antibody repertoire of four *Flavivirus*-experienced donors after their revaccination with the 17DD-YF vaccine. We found that antibody repertoires present common antibody kinetics and dynamics of B cell expansion and diversity after revaccination against YFV. Moreover, we observed a predominance of IgA and IgG antibody lineages under expansion for all individual’s repertoires. On the other hand, we identified unique features of the repertoires that reflect in the magnitude and persistence of the antibody responses over time. Overall, this work sheds light on the complexity of the B cell responses after revaccination against YFV.

## 2. Materials and methods

### 2.1 Human Subjects

Study was approved by the institutional Ethics Committee – Federal University of Minas Gerais, under protocol number CAAE: 08301112.9.0000.5091. Informed consent form and an epidemiological questionnaire were obtained before any sample collection and kept anonymized. Subjects enrolled in this study consisted of three women and one man (n=4), aged between 20 and 45 years old and originally from Belo Horizonte, Minas Gerais – Brazil. All participants were re-vaccinated with YFV-17DD vaccine and a total of 4 ml of venous blood was obtained from each subject before vaccination and on days 7, 14, 28, and 180 following vaccination. Samples were processed by centrifugation at 1,300 rpm and room temperature to separate plasma. Peripheral Blood Mononuclear Cells (PBMC) were also isolated using Ficoll-Paque^TM^ gradient centrifugation and cryopreserved in FBS 90% / DMSO 10%. Both plasma and PBMCs aliquots were stored at -80 °C until further analysis.

### 2.2 Indirect ELISA Assay

Indirect ELISA assay was performed at the Laboratório de Tecnologia Imunológica, Bio-Manguinhos (LATIM, FIOCRUZ-RJ, Brazil) to confirm the presence of IgG antibodies against YFV, DENV1-4, and ZIKV in the plasma of vaccinated subjects. Briefly, 96-well plates were coated with 2.5 μg/ml of whole yellow fever virus particle diluted in carbonate–bicarbonate buffer (pH 9.6) and incubated overnight at 4 °C, as described by Abreu et al., [26]. Then, plates were washed five times with PBS/T (PBS pH 7.4 with 0.05% (v/v) of Tween-20) and blocked with a blocking/diluent solution (BDS) (PBS/T, 0.05% (v/v) of BSA, 3% (w/v) of fetal bovine serum (FBS) and 5% (w/v) of skimmed milk). After 1h incubation at 37 °C, plates were rinsed with PBS/T as previously, and serum samples were serially diluted. An anti-YFV Serum was used to derive a standard curve, in the range of 1 to 0.015 IU/ml. Plates were washed again after 1h of incubation of the samples at room temperature. An anti-human IgG peroxidase conjugated (Sigma) was diluted 1:5000 in BDS and then added to the plates for 1h incubation at RT. After a final washing, 100 μl/well of substrate solution (TMB Plus^TM^ kem-en-tec) were added and after 15 min the reaction was stopped by adding stop-solution (2M H_2_SO_4_). The endpoint measurements were done at 450 nm and the absorbances of the serum sample dilutions were plotted on the standard curve. The results obtained by absorbance values > 0.150 were calculated using the software SoftMax Pro^®^ by regression logistic for four parameters and the antibody titers were expressed in IU/mL.

The detection of Zika and Dengue 1-4 antibodies was carried out using commercially available ELISA kits (Zika-Euroimmun commercial and Panbio dengue IgG indirect ELISA kit, Abbott Laboratories, Chicago, IL, USA) and following the manufacturer’s instructions. Results were expressed as relative units/ml.

### 2.3 Neutralizing Antibody Quantification

Quantification of neutralizing antibody (nAb) against YFV, DENV1-4 and ZIKV in inactivated (56°C/30min.) plasma samples collected from all donors in each time point - 0, 7, 14, 28, and 180 days after YFV-17DD vaccination were performed by the Immunomolecular Analysis Laboratory of Bio-Manguinhos/Fiocruz. Both YF- and DENV-nAb levels were quantified using the micro-focus reduction neutralization test (mFRNT) in 96-well plates, and nAb titers were expressed as the highest serum dilution resulting in 50% focus reduction (mFRNT_50_) [27]. Briefly, samples were serially diluted from 1:3 to 1:729 (dilution factor = 3), followed by the addition of virus suspension (YFV-17D-213/77 vaccine virus; D1-West Pac 74; D2-S16803; D3-CH53489 or D4-341750) containing approximately 100 PFU, resulting in final dilutions of 1:6 to 1:1458 (dilution factor = 2). Plates were incubated at either 37 °C (YFV) or 35 °C (DENV) for 2 hours in a 5% CO_2_ atmosphere for neutralization. The mixtures (diluted samples + virus) were then transferred onto pre-formed confluent Vero CCL-81 cell monolayers for mFRNT-YF or Vero NIBSC cell monolayers, for mFRNT-DENV1-4, both with 2 x 10^5^ cells/well. Carboxymethylcellulose (CMC) was added to each well before final incubation only for mFRNT-YF. The plates were then incubated for 48 hours at either 37 °C (mFRNT-YF) or 35 °C (mFRNT-DENV1-4) with 5% CO_2_. Subsequently, cells were fixed with a solution of ethanol and methanol (1:1), followed by incubation with 4G2 HRP-conjugated monoclonal antibody for 2 hours at 35 °C in a 5% CO_2_ atmosphere. True Blue Dye substrate was added, and after incubation, monolayers were washed, dried, and imaged using an automated acquisition system (28). Focus was manually counted, and samples with titters ≥ 1:100 and ≥ 1:30 were considered seropositive to YFV and DENV, respectively. The cutoff was defined by a ROC curve analysis (data not shown) that compared results of negative and positive samples tested by classical PRNT and microFRNT. This allows the microFRNT results to be correlated with the classical PRNT as previously established by our research group [26, 29–31].

The PRNT_90_-ZIKV (PRNT - plaque reduction neutralization test) was conducted using Vero CCL-81 cells to assess the presence of nAb against ZIKV in samples. Briefly, samples were serially diluted from 1:5 to 1:15,625 (dilution factor = 5), followed by the addition of 60 µL of ZIKV (strain isolated in João Pessoa, Recife State (SisGen A6E5819) suspension containing approximately 100 PFU, resulting in final dilutions of 1:10 to 1:31,250 (dilution factor = 2) [26]. Plates were incubated at 37 °C for 1 hour in a 5% CO_2_ atmosphere for neutralization. For the adsorption step, 100 µL of dilution mixture was added to monolayers in 24-well plates, previously prepared with 2 x 10^5^ cells/well, and incubated for 1 hour at 37 °C with 5% CO_2_. Subsequently, supernatants were discarded, and monolayers were overlaid with semi-solid medium (sE199 plus 2% CMC), then incubated for 4 days at 37 °C in 5% CO_2_. Cell monolayers were fixed with formalin solution (5%) and stained with crystal violet for plaque counting. Plates were photographed using BioSpot^®^ Software Suite (CTL - Cellular Technology Limited, USA), and plaques were manually counted. nAb titers were expressed as the highest serum dilution that resulted in 90% plaque reduction, with samples exhibiting titers ≥ 1:140 were considered seropositive to ZIKV, since the samples were positive for other *Flavivirus* [26].

### 2.4 Antibody BCR Repertoire Sequencing

#### 2.4.1 RNA Extraction

To perform the antibody BCR repertoire analysis, frozen PBMC obtained from each donor before (T0) and on days 7 and 14 after vaccination, were briefly thawed in water bath at 37 °C degree and kept on ice. Samples were resuspended in 1 ml of Trizol and incubated at room temperature for 5 minutes, after washing them in 10 ml of DPBS by centrifugation (500 x g for 10 minutes at 4 °C). Cells were reincubated at the same conditions following addition of 200 µl of chloroform and vortex homogenization. Afterwards, samples were centrifuged at 12,000 x g for 15 minutes at 4 °C to form a three-phase solution.

The aqueous phase and an equivalent volume of isopropanol were transferred to 1.5 ml eppendorf. Samples were gently homogenized by inversion and incubated overnight at -20 °C. The RNA was obtained by centrifuging samples in the next day at 12,000 x g for 15 minutes at 4 °C. The supernatant was discarded and the tube containing the RNA pellet was resuspended in 1ml of 70% ethanol to be washed by centrifugation at 12,000 x g for 10 minutes at 4 °C. After one more wash step, the supernatant was discarded and the remaining ethanol was left to evaporate from the RNA pellet at room temperature. Finally, samples were resuspended in 30 µl of DEPC water and the RNA concentration was measured by Nanodrop and Qubit RNA BR Assay kit (Thermo Fisher Scientific). The extracted RNA was frozen at -80 °C until use.

#### 2.4.2 Isolation and Amplification of VH Antibody BCR Transcripts

The resulting RNA (500 ng) was used for first strand cDNA synthesis using the SuperScript IV enzyme (Thermo Fisher Scientific). IGVH amplification of IgG, IgA and IgM were amplified using the set of primers listed at Table 1 in two different combinations: firstly, 0.2 mM of forward primers (1 to 8; Table 1) were used with 0.1 mM of each IgG and IgA reverse primers (9 and 10; Table 1). Secondly, 0.2 mM of forward primers (1 to 8; Table 1) were used with 0.2 mM of IgM reverse primers (11; Table 1). All reactions were prepared using 0.2 units (U) of Platinum™ Taq DNA Polymerase High Fidelity (Invitrogen), 1X High Fidelity PCR Buffer solution, 0.2 mM of dNTPs, 1 mM of MgSO_4_, and 2 µL of cDNA in a final volume of 50 µL. The PCR reaction was conducted using the following conditions: 2 min at 95 °C; four cycles of 94 °C for 30 s, 50 °C for 30 s, 68 °C for 1 min; four cycles of 94 °C for 30 s, 55 °C for 30 s, 68 °C for 1 min; 22 cycles of 94 °C for 30 s, 63 °C for 30 s, 68 °C for 1 min; 68 °C for 7 min; hold at 4 °C. Products were analyzed on 1% agarose gel stained with Sybr Safe (Invitrogen). Bands (450 bp) were excised and purified with PCR clean-up Gel extraction (NucleoSpin - Macherey-Nagel) following the manufacterer’s instructions. Purified DNA was quantified by Nanodrop and Qubit DNA High Sensitivity kit (Thermo Fisher Scientific).

**Table 1.**
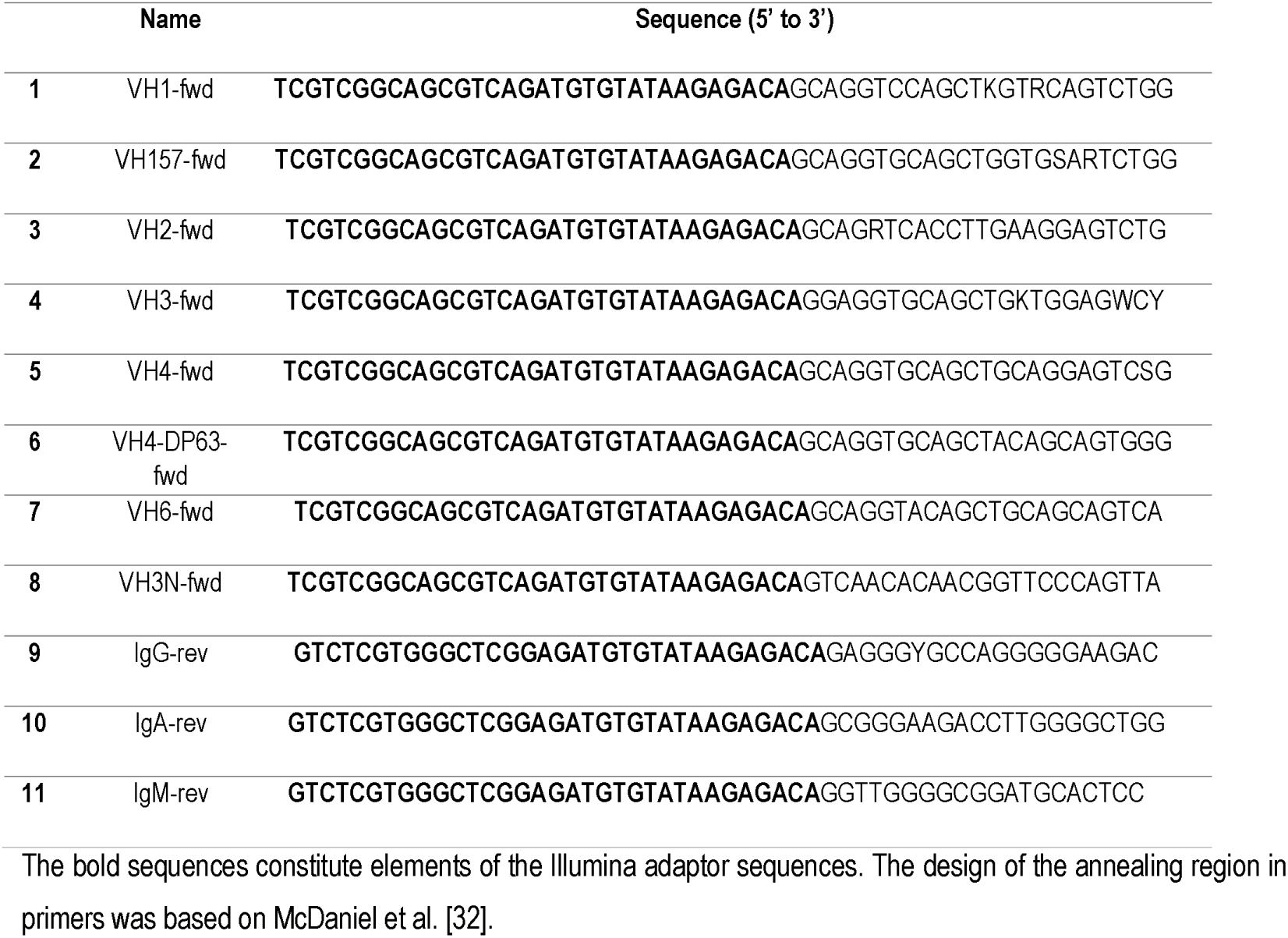
Primer sequences employed for the amplification of antibody-heavy chains (IGVH)

#### 2.4.3 Library Preparation and Sequencing

Sequencing libraries were constructed with 50 ng of each amplicon and using the Nextera XT DNA Library Prep Kit (Illumina) according to the manufacturer’s instructions. The P5 and P7 indexes and adapters were incorporated into the 500 bp amplicons by the overhang adapters added to the primers. All reactions were set in a total of 50 µl under the following conditions: 3 min at 95 °C; 12 cycles of 94 °C for 30 s, 55 °C for 30 s and 68 °C for 1 min; 68°C for 5 min; and hold at 4 °C. Products were purified using of Agencourt AMPure XP beads and the final concentration of the library was quantified using the Qubit DNA High Sensitivity kit (ThermoFisher Scientific). The size and quality of the amplicons were confirmed using the High Sensitivity DNA Kit (Agilent) run on the Bioanalyzer 2100 (Agilent). The VH amplicon library containing the indexes with the final concentration of 18 pM of IgM and 18pM of IgG/IgA samples from the four individuals repertoires were submitted for sequencing in 2×300 paired-end Illumina MiSeq platform.

### 2.5 Bioinformatic Analysis

#### 2.5.1 Raw Reads Processing and Sequence Annotation

Raw Illumina MiSeq output reads were pre-processed and annotated using pRESTO [33] and Change-O [34] packages from Immcantation framework, Biopython [35], and IgBlast [36]. 3’-5’ and 5’-3’ raw reads were joined using the pRESTO AssemblePairs module, with a minimum overlap of 50 nucleotides. The generated sequences were quality filtered using the Biopython Bio.SeqIO.QualityIO module, retaining those with average phred score ≥30.

The pRESTO MaskPrimers module was employed to find on sequences the primers of the IGHV gene and constant region (Table 1, without Illumina adapters). The option score was used, so sequences were kept intact. If IGHV gene or constant region primers were not found, the sequence was filtered out. Sequence annotation was performed with the AssignGenes module from Change-O together with IgBlast to verify similarities between sequences and human germline genes. IMGT human germline reference database was used as reference for V, J, and D gene, as well as their numbering system [37]. After that, Change-O MakeDB module was run to organize the annotation into tabular .tsv files in the MiAIRR format [38]. These files contain information of V, D, J genes and alleles, framework (FW), complementary regions (CDR) sequences, and functional status of the sequence (productive or unproductive). For each PBMC sample (four donors, three times each), it was generated two files, one containing IgM annotated productive sequences and other containing IgG and IgA annotated productive sequences. Based on previously annotated constant region primers, IgA and IgG productive sequences, that were amplified and sequenced together, were splitted into isotype files. Thus, three files from each sample (donor/time) were used in the following analysis. Each one of these files corresponds to a repertoire isotype.

#### 2.5.2 Lineage Definition and Frequency Calculation

To perform lineage analyses, all productive sequences from all samples with the same VJ segment, CDRH3 length and having approximately 90% similarity were grouped into clonotypes using the tool YClon [39]. The default parameters were used for clonotyping, generating clusters with 91% of similarity between the nucleotide sequences. In order to calculate the frequency of a lineage in an isotype repertoire (IgM, IgG or IgA), the number of sequences grouped in this lineage was divided by the total number of sequences that belongs to this isotype repertoire and multiplied by 100. The level of expansion of a lineage was determined by this frequency. If a lineage represented 0.1% or more of an isotype repertoire, it was considered expanded.

#### 2.5.3 Persisting-Expanded and New Expanded Lineages Definition

To assess B cell recall responses following booster YFV vaccination, we analyzed each isotype repertoire and classified expanded B cell lineages (≥ 0.1% frequency) into two groups. Persisting-expanded lineages were those present at baseline (T0), that persisted at T7 and/or T14, and showed expansion post-vaccination. New expanded lineages were those not detected at T0 but emerged and expanded at T7 and/or T14 following vaccination.

To determine the proportion of persisting-expanded and new expanded lineages among the most dominant clones at T14, we first ranked all expanded lineages (≥ 0.1% frequency) from the IgG, IgM, and IgA repertoires in descending order based on their frequency. The top 50 most expanded lineages were then selected and classified as either persisting-expanded or new expanded.

#### 2.5.4 Repertoire Characterization

Repertoire parameters, such as diversity, clonal expansion, size, gene segment usage, somatic hypermutation frequency, and composition and amino acid groups of CDRH3, were analyzed in R studio. Diversity was defined by the Shannon-Wiener index calculated as in McCune et al. [40]. Gene segments and gene subgroups were defined by taking the first assigned gene segment to each sequence from the IgBlast output. After that it was possible to evaluate the frequencies of gene segment usage, gene subgroups, and the combination of V(D)J genes in each repertoire. Somatic hypermutation frequency was defined as one minus the v gene identity assigned by IgBlast, multiplied by 100. R studio was also used to generate the clonal expansion diagram that considered the clonal expansion frequency as a measure to set the expansion state of the IgG, IgA and IgM repertoires.

#### 2.5.5 Comparative Analysis with Known Antibodies

We used the “search by keyword” function of The Patent and Literature Antibody Database (PLAbDab) [41] available in July 2024 with "DENV|Dengue|ZIKV|Zika|YFV|Yellow Fever" expression to acquire a list of known antibodies that bind to DENV, ZIKV, and YFV. To this list, we added the antibodies obtained from Wec et al [12]. To verify if any CDRH3 from this list had at least 80% amino acid identity to any expanded sequence, we translated the nucleotide sequence of the heavy chain sequences from the list and renumbered them using ANARCI [42] to standardize the CDRH3 region definition. Following that, we used PairwiseAligner module from Biopython [35] to make a pairwise alignment of each CDRH3 sequences from the literature list with each sequence from the repertoire, counting the number of amino acid matches. If this count, divided by the larger CDRH3 from the analyzed pair of sequences, is equal or greater than 80%, we would identify them as having similar enough CDRH3. In order to get epitope information of each literature match, we consulted the reference hits’ patent and/or pdb registration according to data availability.

### 2.6 Statistical Analysis

Statistical analysis were conducted in GraphPad Prism version 9.0.0 (GraphPad Software Inc., La Jolla, CA, USA). Shapiro-Wilk test was performed to determine the normality of the data, followed by Mann-Whitney test. Non-parametric Mann–Whitney U tests and analysis of variance tests (anova and Kruskal–Wallis H tests) were used to determine the statistical significance.The mean of each donor-specific paramenter, followed by the average mean of all four donors, as well as the standard deviation were used to compare antibody repertoires differences.

## 3. RESULTS

### 3.1 17DD-YF Booster Induces a Transient Increase in YFV-Neutralizing Antibodies with Limited Impact on DENV and ZIKV Neutralization

Healthy donors (n=4) from a region with high YFV incidence received a standard booster dose of the 17DD-YF vaccine. All participants reported prior YFV vaccination over 10 years ago (Table 2). Blood samples were collected before vaccination (T0) and at days 7, 14, 28, and 180 post-vaccination (Fig. 1A).

**Figure 1:**
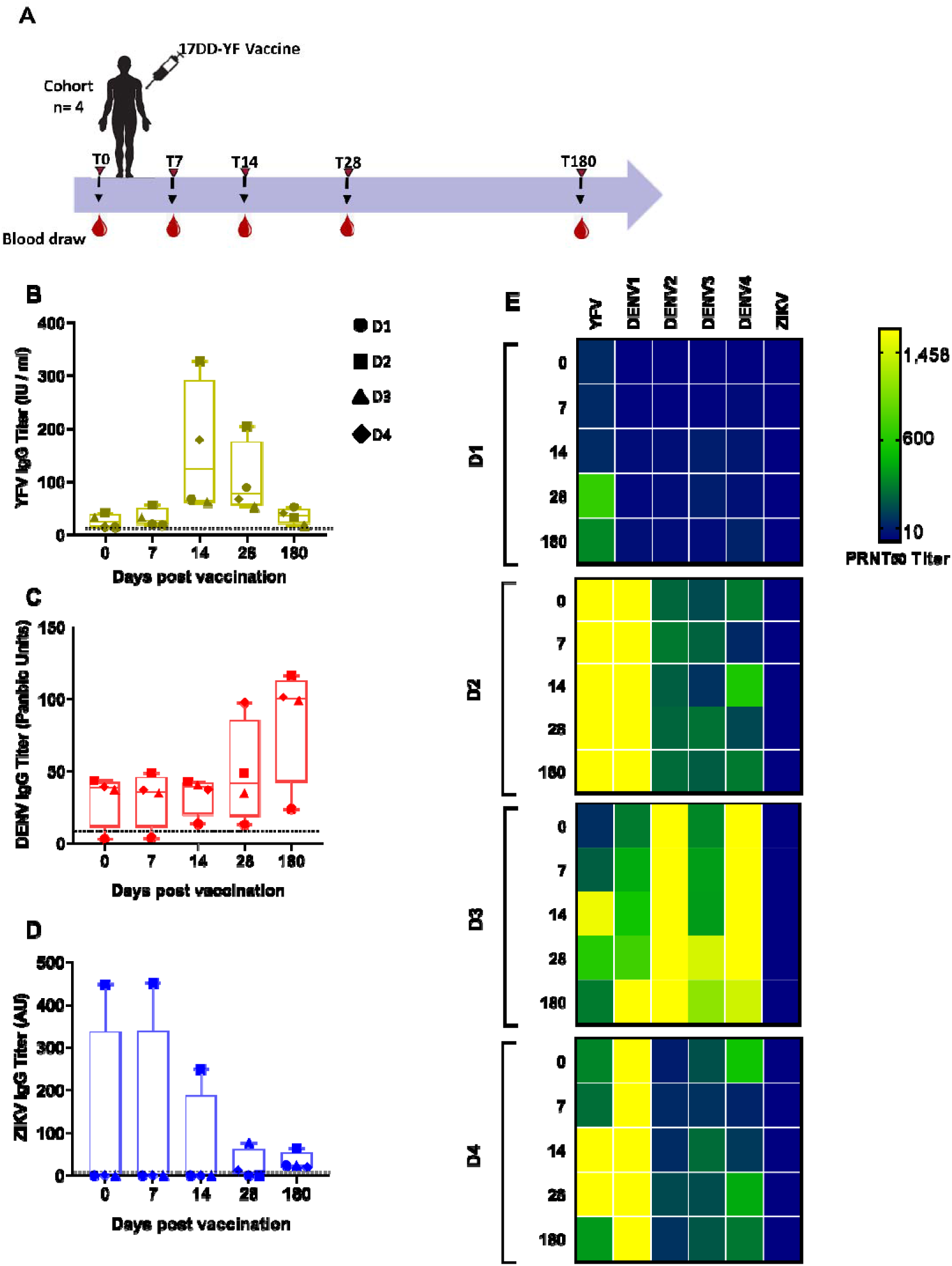
Longitudinal flavivirus antibody responses. **A** Schematic representation of YFV vaccination and longitudinal blood sampling. 17DD-YF vaccine was used to immunize four healthy donors against yellow fever virus. Blood draws were performed before (T0), 7, 14, 28 and 180 days following vaccination to obtain both plasma and PBMCs from each donor at each time point. **B-D** IgG responses against YFV (B), DENV (C) and ZIKV (D). Binding levels were measured using indirect ELISA and the data was expressed as mean IU/ml for YFV, Panbio Units for DENV and AU for ZIKV. Dot lines represent the cut-off for positivity. **E** Neutralizing antibody titers against YFV, DENV1-4, and ZIKV presented by each donor at distinct time points. Results were obtained by performing soroneutralization assay and cut-off criterion for positive mFRNT results were: ≥ 1:100 (YFV), ≥ 1:30 (DENV1-4), and PRNT results ≥1:140 (ZIKV).

**Table 2.** Overview of the study participants and their baseline immune status.

| Total n=4 |  |  |  |  |  |  |  |  |  |
| --- | --- | --- | --- | --- | --- | --- | --- | --- | --- |
| Donor | Sex | Age | Past YFV vaccination | YFV | DENV1 | DENV2 | DENV3 | DENV4 | ZIKV |
| mFRNT<sub>50</sub>/PRNT<sub>90</sub> Seroactivity |  |  |  |  |  |  |  |  |  |
| D1 | F | 38 | Yes | -- | -- | -- | -- | -- | -- |
| D2 | M | 23 | Yes | ++ | ++ | ++ | ++ | ++ | -- |
| D3 | F | 24 | Yes | -- | ++ | ++ | ++ | ++ | -- |
| D4 | F | 41 | Yes | ++ | ++ | -- | ++ | ++ | -- |

F; female. M; male. --; negative. ++; positive. Soroneutralization cut-off for positive mFRNT<sub>50</sub> results: $\geq 1:100$ (YFV), $\geq 1:30$ (DENV1-4); PRNT<sub>90</sub> $\geq 1:140$ (ZIKV).

Polyclonal sera collected at all five time points were analyzed by indirect ELISA against YFV, DENV, and ZIKV to assess the longitudinal dynamics of the antibody response to YFV revaccination and potential cross-reactivity with other flaviviruses. Overall, the majority of donors (50% to 75%) exhibited detectable levels of YFV and/or DENV-specific antibodies at baseline (T0), while only donor 2 displayed high titers of anti-ZIKV IgG prior to immunization (Table 2, Fig. 1B-C). Interestingly, this donor showed a transient decline in anti-ZIKV IgG levels post-vaccination, eventually reaching titers comparable to the other individuals by day 180 (Fig. 1D). In contrast, all donors exhibited increased anti-DENV IgG titers following vaccination, peaking at day 180 (Fig. 1C). Anti-YFV IgG responses followed a consistent pattern among donors, with titers peaking at day 14 and gradually declining to near-baseline levels over six months (Fig. 1B).

To determine whether these binding antibody responses were associated with functional neutralizing activity, we performed neutralization assays (mFRNT50 for YFV and DENV, and PRNT90 for ZIKV). These assays were conducted against YFV, ZIKV, and DENV serotypes 1–4. In line with the ELISA data, three donors exhibited an increase in YFV neutralizing antibody (nAb) titers by days 14 and 28 post-vaccination, followed by a decline by day 180 (Fig. 1E). Notably, there was considerable inter-individual variation in the magnitude of anti-YFV neutralizing responses, likely reflecting differences in baseline immune status (Fig. 1E, Table 2). By contrast, ZIKV neutralization was undetectable across all time points. All donors displayed varying degrees of DENV1-4 neutralization; however, these responses did not parallel the increase in binding titers observed by ELISA. While ELISA indicated enhanced DENV reactivity post-vaccination, DENV nAb titers remained largely unchanged, except for donor 3, who already had detectable DENV nAbs at baseline and showed an increase in nAbs specifically against DENV-1 and DENV-3 following revaccination (Fig. 1E).

Taken together, these findings indicate pre-existing immunity to YFV and/or DENV in the studied individuals, with immune profiles that were modulated—but not uniformly boosted— by YFV revaccination. ZIKV-specific nAbs were not induced, and no evidence of the increase in the cross-reactivity was observed following vaccination. Although YFV-specific binding and neutralizing antibody titers were transiently boosted, both declined markedly within six months post-immunization. In contrast, no consistent pattern of increase or decay was observed for DENV or ZIKV neutralizing antibodies following the booster dose.

### 3.2 YFV revaccination induces a marked expansion of the antibody repertoire by day 14, predominantly involving clonally expanded IgA and IgG lineages

During revaccination, both naïve and memory B cells (MBCs) are activated, leading to the production of antibodies with distinct specificities. Analyzing B cell repertoire dynamics following vaccination has been shown to predict the nature and magnitude of antibody responses [43]. In the case of yellow fever virus (YFV) vaccination, plasmablast responses typically peak between 10 and 14 days post-vaccination [12]. Based on this, we sequenced the VH region from bulk PBMCs collected on days 0, 7, and 14 after booster immunization to investigate the temporal dynamics of the antibody repertoire.

We analyzed the immunoglobulin heavy chain (IgH) repertoires of IgG, IgM, and IgA isotypes, including both the variable (VDJ exon) and constant regions, to monitor repertoire composition and clonal changes over time. On average, 69,254 B cell lineages were analyzed across all samples (Table S1).

A small fraction of B cell lineages (0.28% to 1.96%) was shared across time points within each isotype, indicating the persistence of specific clones and suggesting a recall response to the booster dose (Fig. 2A). Because antigen-driven clonal expansion typically reduces repertoire diversity, we next evaluated how the YFV booster impacted diversity and expansion within the first two weeks post-vaccination.

**Figure 2:**
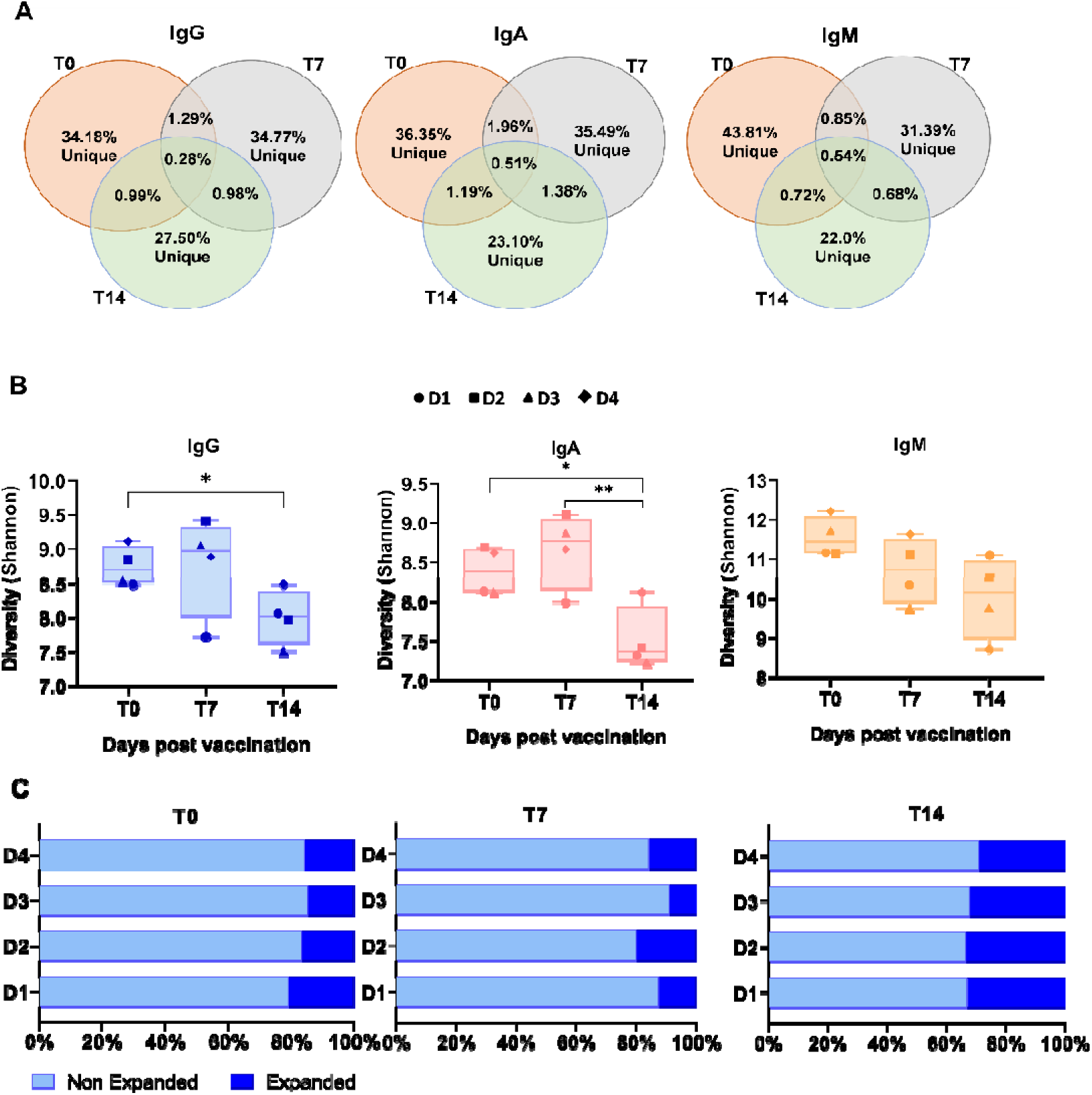
Dynamics of repertoire diversity and expansion following 17DD-YF vaccine boost. **A** Average of IgG, IgA and IgM lineages across donors that are and are not shared between the indicated time points 0, 7 and 14 days after vaccination. **B** Diversity of IgG (B), IgA (C) and IgM (D) repertoires was assessed by Shannon diversity index at 0, 7 and 14 days following vaccination. **C** Proportion of expanded and non expanded IgG, IgA and IgM lineages in each vaccinee repertoire at 0, 7 and 14 days after 17DD-YF booster dose. Lineages were classified into non expanded (< 0.1%) and expanded (≥ 0.1%) according to their frequencies in the repertoire. \**p* < 0.01 and ** *p* ≤ 0.005.

All donors exhibited a substantial reduction in repertoire diversity by day 14 across all isotypes (Fig. 2B), coinciding with increased clonal expansion. Notably, expanded lineages (≥ 0.1% frequency) represented approximately 30% of the repertoire at this time point (Fig. 2C). Isotype-specific analysis of the top 200 lineages revealed a predominant expansion of IgA lineages, followed by IgG, with both peaking at day 14 post-revaccination (Fig. 3).

**Figure 3:**
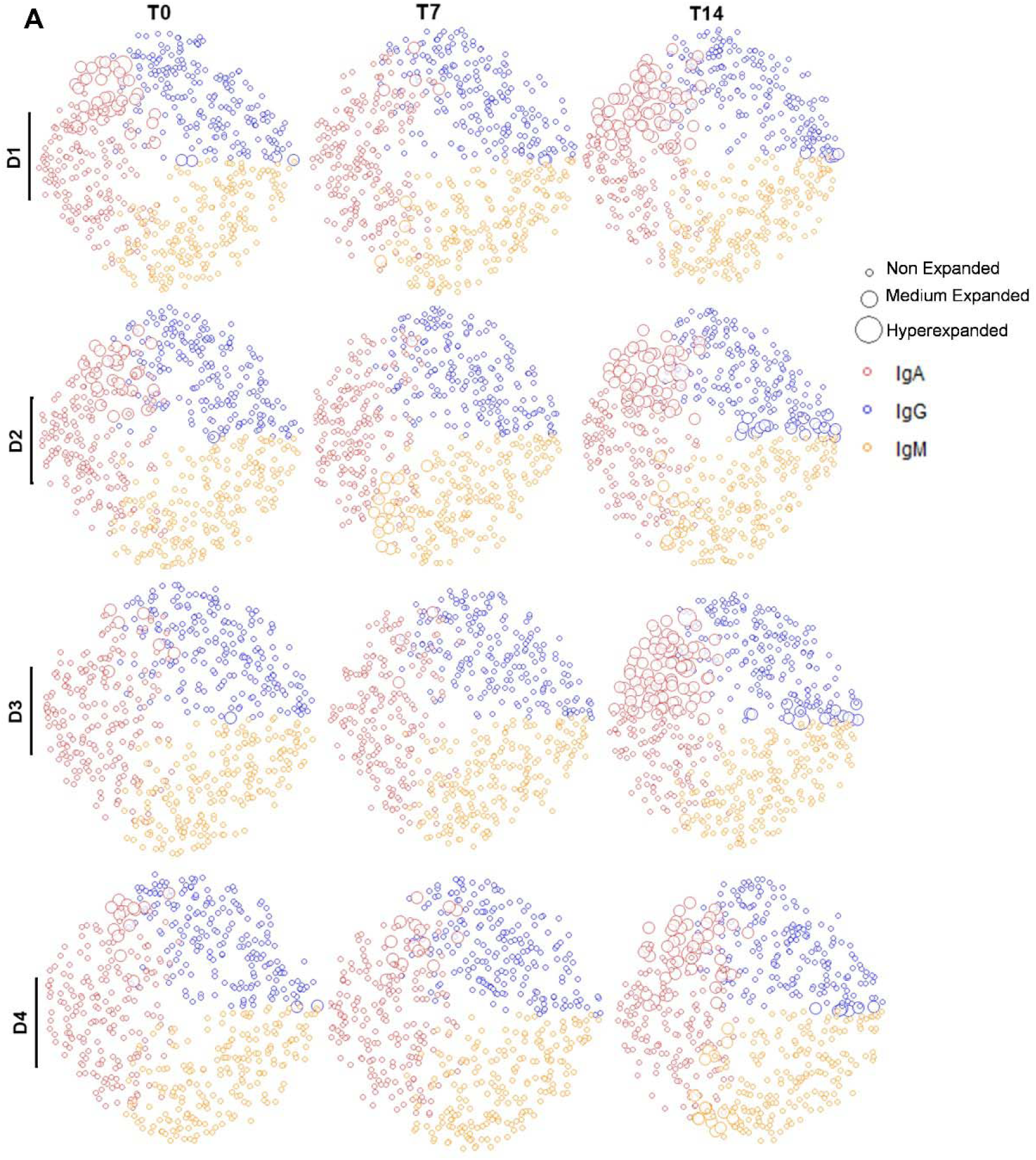
Overview of the repertoire expansion state over the two weeks after 17DD-YF boosting. Diagram representing the top 200 most frequent B cell lineages from each IgG, IgA and IgM repertoire. Each circle represents a B cell lineage with size proportional to its frequency in the repertoires at 0, 7 and 14 days after revaccination. Expansion levels were defined as non expanded for lineages presenting frequency below to 0.1%, medium expanded for lineages with frequency between 0.1% and 1% and hyperexpanded for lineages presenting frequencies greater than 1%.

### 3.3 Low-Frequency Persisting Lineages Dominate Top Expanded B Cell Clones at Day 14 Post-Vaccination

To explore in detail the dynamics of clonal expansion in the B cell repertoire following revaccination, we tracked individual clonal lineages over time to assess whether specific lineages were recalled and maintained within the repertoire. Previous analyses had already revealed the presence of shared lineages across time points (Fig. 2A). To determine whether these persistent B cells were associated with the vaccine response, we defined a “persisting-expanded” lineage as one that was present at baseline (T0), persisted at T7 and/or T14, and showed expansion (≥ 0.1% frequency) after vaccination. Remarkably, less than 30% of all expanded lineages met this criterion—representing an average of only 28 out of 1,003 total lineages (Fig. 4A). The majority of these persisting-expanded lineages were classified as IgA, while IgG and IgM lineages showed lower levels of persistence (Fig. 4B).

**Figure 4:**
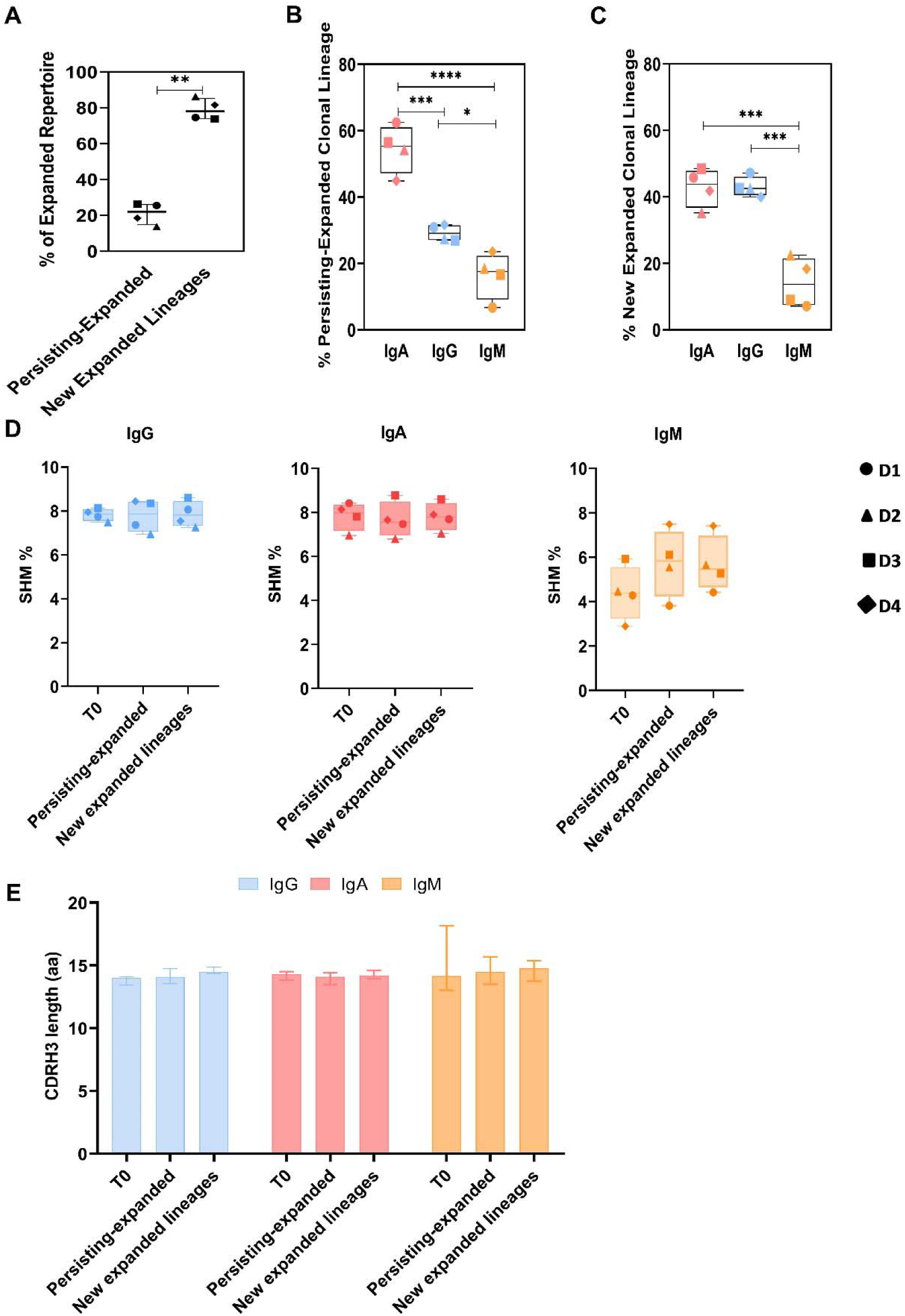
Analysis of persisting-expanded and new lineages. **A** Proportion of persisting-expanded and new expanded lineages found in the repertoires on day 7 and 14. **B-C** Proportion of persisting-expanded and new expanded lineages according to the isotype. **D** SHM average loads presented by IgG, IgA and IgM clonal lineages found in expanded repertoire on day 0 and on T14 by persisting-expanded and new expanded lineages. **E** CDRH3 average length of clonal lineages. \**p* < 0.05, \*\*\**p* ≤ 0.001, \*\*\*\**p* ≤ 0.0001.

Thus, the expanded repertoire was predominantly (>70%) composed of lineages detected exclusively after the booster vaccination (T7 and/or T14), with an average of 58 such lineages per donor, totaling 2,048 across all donors (Fig. 4A). Although the possibility of sampling bias may partially account for the apparent absence of these lineages at earlier time points, we observed consistent proportions of both persisting and newly emerging lineages across all donors. This pattern suggests a shared and coordinated immune response to the YFV booster vaccination (Fig. 4A). Notably, most of these newly expanded lineages—whether persistent or newly emergent—were of the IgA and IgG isotypes (Fig. 4B–C), aligning with prior observations of the dominant involvement of IgA and IgG in the expanded repertoire (Fig. 3).

To gain insights into the nature of these clonal lineages and their relevance to vaccine-induced immunity, we next analyzed their molecular features. Both persisting-expanded and newly expanded lineages exhibited similar levels of somatic hypermutation (SHM), with median values ranging from 6% to 8%, and comparable CDRH3 lengths (approximately 15 amino acids), with no significant differences observed compared to the pre-vaccination time point (Fig. 4D–E). Interestingly, donor 1—who lacked pre-existing immunity to flaviviruses—had an IgM repertoire with notably lower SHM levels.

We also assessed germline VH gene usage across expanded lineages to determine whether any preferential gene segment convergence indicative of antigen-driven selection was present. Despite similar SHM levels and CDRH3 lengths, we observed non-uniform usage of a broad range of V gene segments in both persisting and new expanded populations (Fig. S.1). Moreover, the VH gene usage profiles at baseline (T0) were similar between these groups, indicating no strong selection for specific VH genes following YFV revaccination.

Given that overall repertoire diversity decreased by day 14 across all isotypes (Fig. 2B), we further examined the 50 most expanded clonal lineages at this time point to assess the contribution of persisting-expanded and newly expanded clones. We found that persisting-expanded lineages accounted for a substantial proportion of the top 50 clones in most donors, with the exception of donor 3 (Fig. 5). These findings indicate that while persisting-expanded lineages represent fewer than 30% of the expanded repertoire overall, they make a significant contribution to clonal dominance, being highly represented among the most abundant lineages.

**Figure 5:**
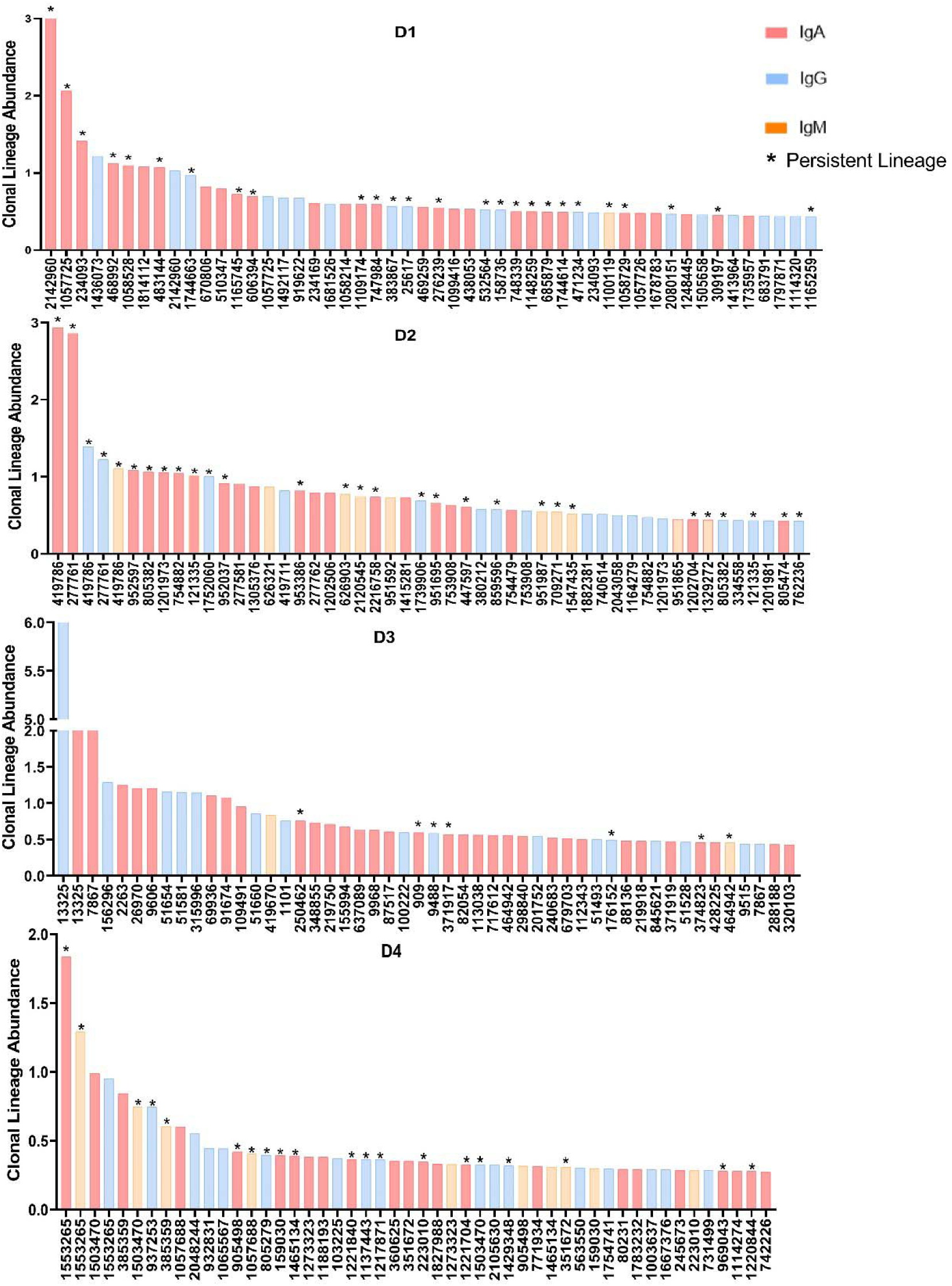
Representative histogram of the top 50 most expanded clonal lineages (IgG, IgA and IgM) that comprise the YFV vaccinees’ repertoire at 14 days after vaccination. Bars indicate the relative frequency of each lineage in the repertoire and * shows the persistent lineages.

### 3.4 B cell repertoires from YFV vaccinees contain lineages matching known anti-YFV, DENV, and ZIKV antibodies

To evaluate the potential cross-reactivity of vaccine-induced B cell lineages, we compared BCR repertoire sequences against a reference database of monoclonal antibodies (mAbs) with experimentally confirmed binding to YFV, DENV, and ZIKV. Specifically, we queried the CDRH3 regions of both expanded and non-expanded IgG, IgA, and IgM lineages from each individual against the antibody sequence database [12, 41]. Alignments with >80% sequence identity were considered matches, providing insights into the identity and potential cross-reactivity of lineages following YFV revaccination.

In total, 57,626 unique CDRH3 sequences matched characterized *Flavivirus* antibody sequences in the database. These matches were grouped into 17,425 unique lineages, of which 27 were expanded and 17,398 were non-expanded post-vaccination (Table S2). The predominance of non-expanded matches suggests that the vaccinees’ repertoires contain DENV- and/or ZIKV-specific B cell lineages that did not expand upon YFV booster vaccination, according to our criteria (clonal frequency >0.1%).

Among the 27 expanded matching lineages, 11 were IgG, 13 IgA, and 3 IgM. Most of these had previously been described as DENV-specific (Fig. 6A, Table S2). A small subset of these matches displayed cross-reactivity to all three flaviviruses—YFV, DENV, and ZIKV (Fig. 6A). To determine whether these cross-reactive clones originated from persisting-expanded or newly expanded lineages, we tracked the origin of each matched clone and found that most came from newly expanded lineages across all isotypes (Fig. 6B).

**Figure 6:**
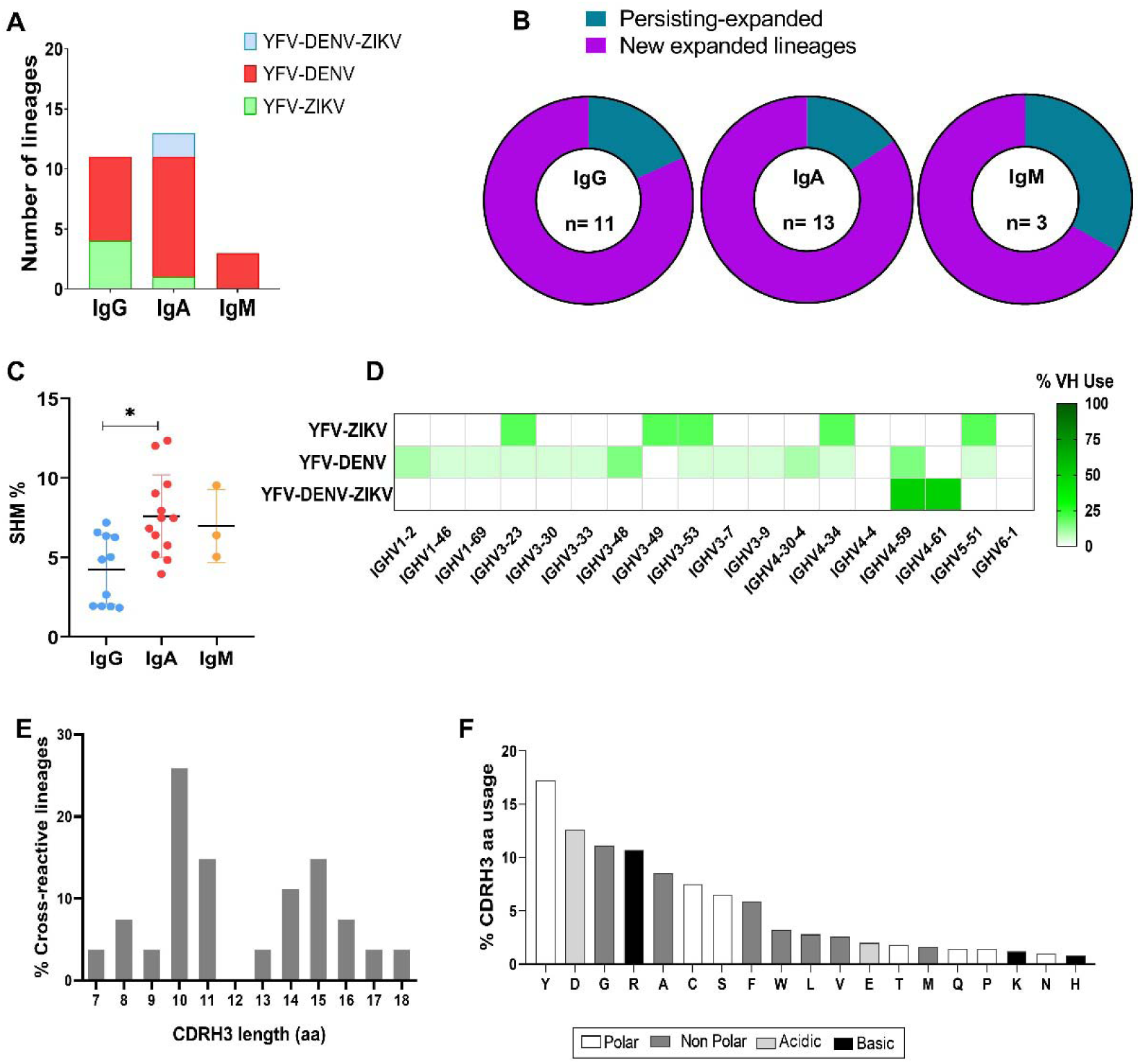
Analysis of putative anti-flaviviruses antibodies present in the analyzed repertoires. **A** Number of discovered hits that display specificity to YFV and cross-reactivity to ZIKV, DENV or ZIKV and DENV. **B** Fraction of cross-reactive lineages that are either persisting-expanded or new expanded lineages. The number in the center of the pie chart shows the total number of lineages belonging to the isotype. **C** SHM rates of IgG, IgA and IgM lineages that may display cross-reactivity. **D** VH germline gene usage of cross-reactivity lineages. **E** CDRH3 length distribution of lineages that may present cross-reactivity to DENV and/or ZIKV. **F** Physicochemical properties and amino acid usage of the cross-reactive lineages CDRH3.

To explore molecular features indicative of potential cross-reactivity, we analyzed SHM levels, VH gene usage, CDRH3 length, and amino acid composition of the expanded matched lineages. SHM levels varied across isotypes, with average mutation rates of 4.5% for IgG, 7.5% for IgA, and 6% for IgM (Fig. 6C). These values are generally consistent with reported SHM levels in healthy individuals: 2.83 ± 0.23% (IgM), 7.24 ± 0.07% (IgG), and 8.37 ± 0.23% (IgA) (44).

Regarding VH gene usage, the cross-reactive expanded lineages utilized a restricted set of germline segments (Fig. 6D), suggesting potential convergence in response to antigenic stimulation. The CDRH3 length distribution in these clones ranged from 7 to 18 amino acids, with most IgG, IgA, and IgM lineages showing a 10-amino-acid CDRH3 (Fig. 6E).

In terms of physicochemical properties, the CDRH3 regions of cross-reactive clones were predominantly hydrophilic, characterized by a high content of polar amino acids such as tyrosine, cysteine, and serine (Fig. 6F).

Taken together, these findings demonstrate that the post-revaccination repertoire contains B cell lineages with sequence similarity to known DENV- and/or ZIKV-reactive antibodies, some of which expand following YF-17DD boosting. Moreover, these lineages often display signs of affinity maturation, which may contribute to effective recognition and potential cross-neutralization of flaviviral antigens.

### 3.5 Most literature-matched anti-*Flavivirus* antibodies in the YFV vaccinees’ expanded repertoire likely target the DIII domain and fusion loop of the envelope protein

We aimed to identify the epitopes targeted by the matched B cell lineages and assess whether there was evidence of an immunodominance hierarchy among them. To achieve this, we analyzed the experimentally characterized monoclonal antibodies (mAbs) from the literature that comprise the database used in our cross-reactivity analysis. Based on these known mAbs, we identified two structural proteins most frequently recognized: the envelope (E) protein and the precursor membrane protein (prM) (Fig. 7A).

**Figure 7:**
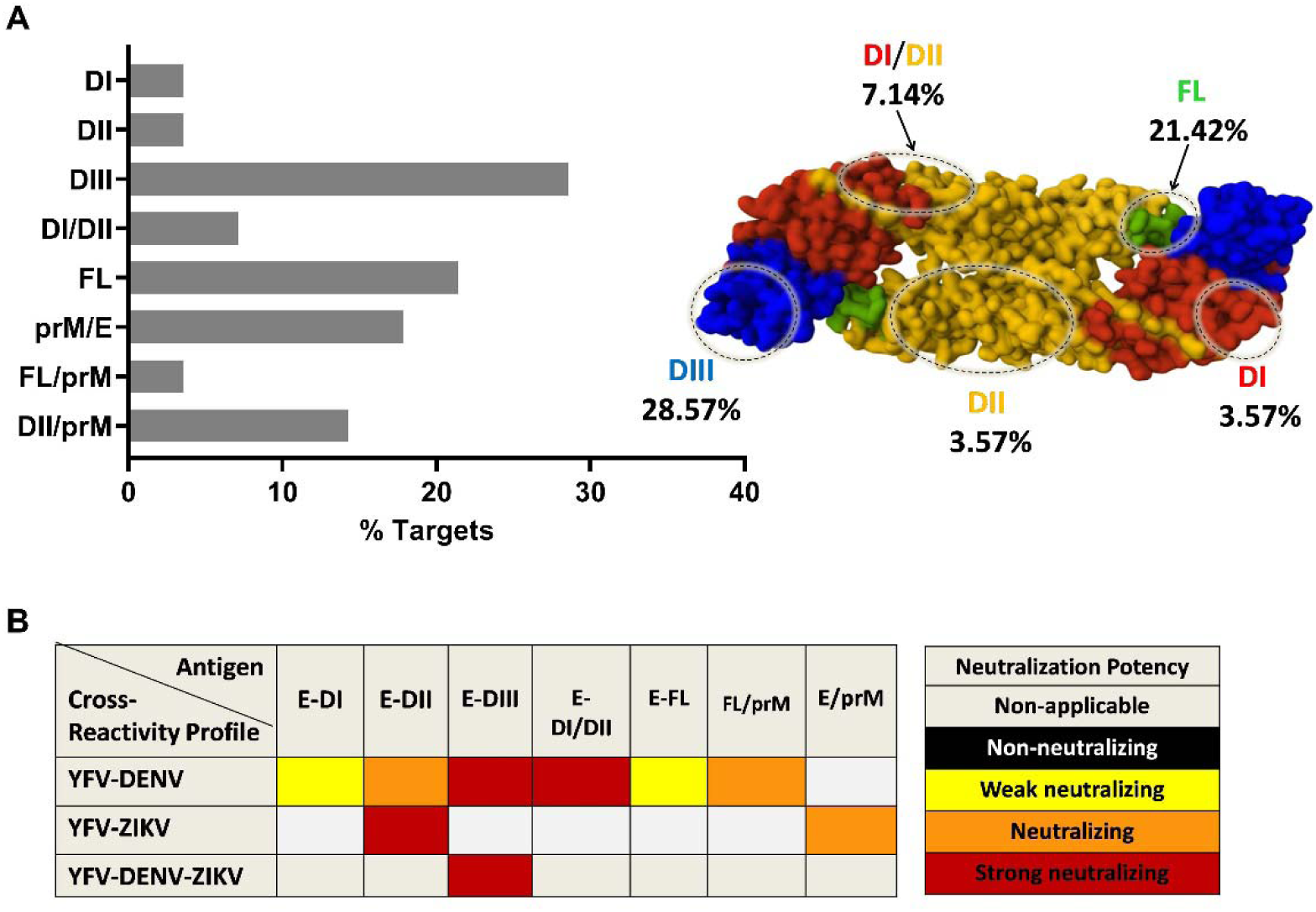
Antigenic sites and functional properties of the lineages elicited by 17DD-YF vaccine booster. **A** Antigenic sites targeted by cross-reactive lineages. Crystal structure of YFV E dimer (Protein Data Bank ID code 6IW2) represented as space filling model. Domains I, II and III are highlighted in red, yellow and blue respectively, in each monomer. FL is colored in green. Labels show the proportion of the mAbs targeting epitopes within each structural region. **B** Prediction of neutralization potential of each mAb according to data in published literature.

Consistent with published findings, the majority of these mAbs target the E protein. To further pinpoint the specific epitope regions within E, we examined their structural localization and found that most of the matched mAbs bind to the domain III (DIII) of the E protein (28.57%) or to the fusion loop (FL) located at the tip of domain II (DII) (21.42%) (Fig. 7A). A notable proportion of mAbs (17.86%) were found to target epitopes spanning both the E and prM proteins (Fig. 7A).

We next assessed the relationship between epitope location and neutralization potency, based on experimental data reported for these mAbs in the literature. Interestingly, antibodies targeting the FL and domain I (DI) were generally described as weakly neutralizing, whereas those directed toward DIII and quaternary epitopes involving DI and DII were associated with strong neutralization capacity (Fig. 7B).

Although limited, our findings suggest that some of the matched lineages may correspond to cross-reactive antibodies targeting functionally relevant epitopes, such as DIII and quaternary regions of the E protein, with potential for neutralizing activity against ZIKV and DENV.

## 4. Discussion

The global emergence of flaviviruses over the past several decades has threatened millions of people and become a major public health concern worldwide. The lack of specific antiviral treatments has worsened disease management, making prevention through vaccination the primary countermeasure for some viruses, such as YFV. A detailed characterization of B cell repertoire dynamics and kinetics following YFV vaccination can provide important insights into long-term protection and inform the necessity of booster campaigns to maintain population immunity.

In this study, we analyzed the antibody repertoire of four healthy individuals previously vaccinated against YFV, all originating from an endemic region with co-circulation of DENV, ZIKV, and YFV. Our findings offer insights into the humoral immune response to 17DD-YF booster vaccination, at both the cellular and serological levels. Although the literature provides extensive data on humoral responses to the 17DD-YF vaccine, particularly in terms of neutralizing antibody titers, no previous studies have investigated the peripheral B cell repertoire after a booster dose and its relation to serological responses.

A key observation from our analysis was the marked clonal expansion occurring 14 days after revaccination, which was accompanied by changes in antibody titers and neutralizing activity. This response pattern mirrors findings from studies examining the primary response to the 17D vaccine [6, 12, 45].

Previous reports have shown that primary 17DD-YF vaccination leads to a peak in antibody levels between days 30 and 45, followed by a gradual decline up to day 365 [30]. Here, we observed that humoral responses following revaccination reached their peak earlier and began to wane by day 28. This is consistent with prior findings indicating that pre-existing immunity may limit viral replication after YFV revaccination, thereby reducing antigen availability and B cell stimulation [46–49]. Interestingly, this kinetic profile was seen in three of the four donors, all of whom exhibited serological evidence of prior exposure to flaviviruses (YFV, DENV, or ZIKV). In contrast, donor 1—who showed no pre-existing immunity to any tested *Flavivirus*—displayed a distinct response pattern. This individual seroconverted to YFV by day 28, and neutralizing antibody titers remained high through day 180. Moreover, the IgM repertoire of this donor exhibited lower SHM levels compared to the others, a result that aligns with antibody dynamics typically seen after primary 17DD-YF vaccination [30].

To our knowledge, this is the first high-throughput sequencing study to characterize the kinetics, diversity, and clonal expansion of the antibody repertoire following a 17DD-YF booster dose. Notably, the early response was dominated by expanded IgA and IgG lineages, with limited involvement of IgM clones. This is consistent with previous reports describing increased IgA plasmablasts following 17D primary vaccination, particularly at day 7, with frequencies peaking around day 14 alongside IgG [6].

Similar observations have been made in primary dengue virus infections, where IgA-expressing plasmablasts dominated the repertoire between days 14 and 16 [50–51]. Although the role of IgA in anti-*Flavivirus* immunity remains unclear, some studies have demonstrated that IgA can neutralize DENV without inducing antibody-dependent enhancement (ADE), due to its low affinity for Fc receptors [52–53]. Furthermore, IgA has been shown to antagonize IgG-mediated ADE by competing for the same epitopes [53]. Taken together, these findings suggest that the presence of IgA antibodies in the YFV vaccinee repertoire may contribute to protective, non-pathogenic immune responses. This is supported by a previous study that found no evidence of increased risk of severe dengue among YFV-vaccinated individuals living in endemic areas [54].

Our data also revealed that a significant portion of pre-existing lineages underwent expansion upon revaccination. Molecular analysis of these persisting-expanded lineages, alongside newly expanded ones, showed similar features in terms of SHM levels, CDRH3 length (∼15 amino acids), and preferential usage of IGHV3 and IGHV4 gene families. Interestingly, Wec et al. [12] reported that primary YFV vaccination in individuals from non-endemic areas resulted in repertoires with low SHM, short CDRH3s (∼10 amino acids), and preferential IGHV3-72 usage by day 14. Similarly, our findings suggest that during secondary vaccination, the repertoire prioritizes the rapid expansion of both pre-existing and newly generated lineages, without extensive affinity maturation.

We also investigated the presence of lineages with potential cross-reactivity to DENV and ZIKV. Using CDRH3 sequence matching against a database of experimentally validated *Flavivirus* antibodies, we identified multiple lineages with high sequence identity. While most of these were not expanded after revaccination—likely reflecting past exposures to DENV and/or ZIKV—some matched lineages did expand following the 17DD-YF booster. Notably, many of these had predicted epitopes in the DIII domain and the fusion loop (FL) of the E protein, consistent with prior characterization of anti-*Flavivirus* antibodies [12, 17–18, 55–57].

In primary YFV vaccinees, most isolated mAbs were non-neutralizing and mapped to the FL [12]. While our number of matched hits was limited—possibly due to the small size of available databases—we could still identify evidence supporting the presence of cross-reactive antibodies in the YFV vaccinees’ repertoire. However, further validation is needed to confirm their abundance, functional activity, and epitope specificity. Importantly, the limited availability of characterized YFV antibodies likely underestimates the extent of cross-reactivity detectable in silico.

Our study has several limitations, including sample size (n=4), the use of bulk PBMCs without single-cell resolution, and the absence of serum antibody data. These factors, along with potential variability in the timing of prior *Flavivirus* exposures, limit our ability to identify all relevant B cell lineages and assess the full impact of revaccination. Additionally, longer follow-up could capture later stages of affinity maturation, which has been shown to continue for months after boosting [12]. Further studies are necessary to integrate cellular and serological analyses for a more complete understanding of the anti-YFV antibody response.

In summary, this study provides a detailed view of the kinetics, magnitude, and composition of the B cell response to 17DD-YF revaccination. Although we acknowledge the need for deeper investigation of antibody abundance and persistence in serum, our work tracks the molecular features of the B cell response and contributes valuable insights into the generation of vaccine-induced immunity. This knowledge may ultimately improve our understanding of vaccine-induced protection, particularly with regard to B cell development, repertoire architecture, and population-level variation in endemic settings.

## CRediT authorship contribution statement

Christina A Martins: Conceptualization, Writing – review & editing, Writing – original draft, Methodology, Investigation, Formal analysis, Data curation. Milene Barbosa Carvalho: Writing – review & editing, Investigation, Methodology, Formal analysis, Data curation. João Diniz Gervásio: Writing – review & editing, Methodology, Formal analysis, Data curation. Carlena Navas: Writing – review & editing, Methodology, Formal analysis, Data curation. Luciana Zuccherato: Methodology, Data curation. Marcele Rocha: Writing – review & editing, Methodology. Manuela Emiliano Ferreira:, Methodology, Data curation. Camila Souza: Methodology. Adriana de Souza Azevedo Soares: Writing – review & editing, Methodology, Formal analysis, Data curation. Brenda de Moura Dias: Methodology, Formal analysis, Data curation. Nathalia dos Santos Alves: Methodology, Formal analysis, Data curation. Sheila Maria Barbosa de Lima: Methodology, Formal analysis, Data curation. Waleska Dias Schwarcz: Methodology, Formal analysis, Data curation. Andréa Marques Vieira da Silva: Methodology, Formal analysis, Data curation. Ana Paula Dinis Ano Bom: Methodology, Formal analysis, Data curation. Camilla Bayma Fernandes: Methodology, Formal analysis, Data curation. Renata Carvalho Pereira: Methodology, Formal analysis, Data curation. Mauro M. Teixeira: Methodology. Jason Lavinder: Writing – review & editing, Resources. Gregory Ippolito: Writing – review & editing, Resources. Liza Figueiredo Felicori: Supervision, Project administration, Investigation, Resources, Funding acquisition, Writing – review & editing, Formal analysis, Data curation, Conceptualization.

## Acknowledgements

We thank Dra. Lísia Esper, and UBS staff for sample collection. We thank Regina Fernandes for the project management and Dr. George Georgiou for comments and his kindly revision to this manuscript. We also thank the volunteers who agreed to participate in this study.

## Data availability

The datasets generated during the current study are deposited in the SRA database under the accession number: PRJNA1150892 (https://www.ncbi.nlm.nih.gov/bioproject/1150892). Additional data not included can be obtained by contacting the corresponding author.

